# A Multi-Epitope/CXCL11 Prime/Pull Coronavirus Mucosal Vaccine Boosts the Frequency and the Function of Lung-Resident CD4^+^ and CD8^+^ Memory T Cells and Protects Against COVID-19-like Symptoms and Death Caused by SARS-CoV-2 infection

**DOI:** 10.1101/2023.05.23.542024

**Authors:** Latifa Zayou, Swayam Prakash, Nisha Rajeswari Dhanushkodi, Afshana Quadiri, Izabela Coimbra Ibraim, Mahmoud Singer, Amirah Salem, Amin Mohammed Shaik, Berfin Suzer, Amruth Chilukuri, Jennifer Tran, Pauline Chau Nguyen, Miyo Sun, Kathy K. Hormi-Carver, Ahmed Belmouden, Hawa Vahed, Jeffrey B. Ulmer, Lbachir BenMohamed

## Abstract

The pandemic of the coronavirus disease 2019 (COVID-19) has created the largest global health crisis in almost a century. Following exposure to SARS-CoV-2, the virus particles replicate in the lungs, induce a “cytokine storm” and potentially cause life-threatening inflammatory disease. Low frequencies of function SARS-CoV-2-specific CD4^+^ and CD8^+^ T cells in the lungs of COVID-19 patients were associated with severe cases of COVID-19. The apparent low level of T cell-attracting CXCL9, CXCL10, and CXCL11 chemokines in infected lungs may not be sufficient enough to assure the sequestration and/or homing of CD4^+^ and CD8^+^ T cells from the circulation into infected lungs. We hypothesize that a Coronavirus vaccine strategy that boosts the frequencies of functional SARS-CoV-2-specific CD4^+^ and CD8^+^ T cells in the lungs would lead to better protection against SARS-CoV-2 infection, COVID19-like symptoms, and death. In the present study, we designed and pre-clinically tested the safety, immunogenicity, and protective efficacy of a novel multi-epitope//CXCL11 prime/pull mucosal Coronavirus vaccine. This prime/pull vaccine strategy consists of intranasal delivery of a lung-tropic adeno-associated virus type 9 (AAV-9) vector that incorporates highly conserved human B, CD4^+^ CD8^+^ cell epitopes of SARS-CoV-2 (*prime*) and pulling the primed B and T cells into the lungs using the T cell attracting chemokine, CXCL-11 (*pull*). We demonstrated that immunization of HLA-DR*0101/HLA-A*0201/hACE2 triple transgenic mice with this multi-epitope//CXCL11 prime/pull Coronavirus mucosal vaccine: (*i*) Increased the frequencies of CD4^+^ and CD8^+^ T_EM_, T_CM_, and T_RM_ cells in the lungs; and (*ii*) reduced COVID19-like symptoms, lowered virus replication, and prevented deaths following challenge with SARS-CoV-2. These findings discuss the importance of bolstering the number and function of lung-resident memory CD4^+^ and CD8^+^ T cells for better protection against SARS-CoV-2 infection, COVID-19-like symptoms, and death.

## INTRODUCTION

Severe acute respiratory syndrome coronavirus 2 (SARS-CoV-2), a novel coronavirus identified at the end of 2019, has led to the current global pandemic. The SARS-CoV-2 virus belongs to the subgenus sarbecovirus of the genus betacoronavirus, the genus from which two SARS-CoV-2 closely related viruses (SARS-CoV-1 and MERS-CoV) have crossed the species barrier to humans over the past two decades (1). SARS-CoV-2 uses angiotensin-converting enzyme 2 (ACE2) receptors to enter and infect pulmonary alveolar cells (2–4). This in turn causes the SARS-CoV-2 virus particles to replicate in the lungs, inducing a phenomenon commonly known as a “cytokine storm” and potentially causing life-threatening inflammatory lung disease (5, 6).

SARS-CoV-specific CD4^+^ and CD8^+^ T cells that reside in the lungs appeared to play a critical role in aborting virus replication. However, low frequencies of functional SARS-CoV-2-specific CD4^+^ and CD8^+^ T cells in the lungs of COVID-19 patients were associated with severe COVID-19 cases (7). Three major SARS-CoV-2-specific memory CD4^+^ and CD8^+^ T cell subsets (i.e. effector memory (T_EM_), resident memory (T_RM_), and central memory (T_CM_)), develop, infiltrate and sequester in the infected lungs in response to three major T-cell-attracting chemokines: CXCL9, CXCL10, and CXCL11 among others and enable induced T cells to relocate to specific sites of infection, such as the lungs (8–10). CXCR3, a common chemokine receptor of CXCL-9, CXCL-10, and CXCL-11, is expressed on effector and memory T cells and is responsible for the chemotaxis of T cells within this axis (11, 12). However, the apparent low levels of T cell-attracting chemokines CXCL9, CXCL10, and CXCL11 in the infected lungs of severely ill COVID-19 patients may not be sufficient enough to assure sequestration of CD4^+^ and CD8^+^ T_RM_ cells or to guide homing CD4^+^ and CD8^+^ T_EM_ and T_CM_ cells from the circulation into infected lungs (12, 13).

Parenteral vaccines, administrated intramuscularly or subcutaneously, are often less effective in boosting the frequencies of functional CD4^+^ and CD8^+^ T cells in mucosal tissues including, the pulmonary mucosal site of SARS-CoV-2 replication (14,15). In contrast, mucosal vaccines can potentially produce both mucosal and systemic T-cell immunity (16). In the present study, we hypothesize that a prime/pull vaccine strategy that consists of (i) First priming SARS-CoV-2-specific B cells, CD4^+^ T cells, and CD8^+^ T cells using the engineered lung-tropic AAV9 vector co-expressing the recently identified immunodominant human CD4^+,^ and CD8^+^ T cell epitopes (designated in this report as CoV-Vaccine) and delivered intranasally followed by; (ii) Pulling the “primed” CD4^+^ T cells, and CD8^+^ T cells into the lungs using CXCL-9, CXCL-10, or CXCL-11 T-cell attracting chemokines delivered intranasally (nose drops), will boost the frequency of functional CD4^+^ and CD8^+^ T cells and led to strong local protective T-cell immunity in the lungs against SARS-CoV-2 infection, COVID19-like symptoms, and death (17).

Using a rational reverse vaccine engineering approach, we designed and pre-clinically tested the safety, immunogenicity, and protective efficacy of three multi-epitope prime/pull Coronavirus mucosal vaccine candidates (18). These Coronavirus mucosal vaccines consist of intranasal delivery of a lung-tropic adeno-associated virus type 9 (AAV-9) vector that incorporates highly conserved human B, CD4^+^ CD8^+^ cell epitopes of SARS-CoV-2 that are selectively recognized by the B and T cells from naturally protected asymptomatic COVID patients (*prime*), and pulling the primed T cells into the mucosal sites of the lungs using the T cell attracting chemokines CXCL-9 CXCL-10 or CXCL-11 (*pull*) (19). We demonstrated that immunization of a novel HLA-DR*0101/HLA-A*0201/hACE2 triple transgenic mouse model with a multi-epitope//CXCL11 prime/pull Coronavirus mucosal vaccine boosted the highest frequencies of functional lung-resident CD4^+^ and CD8^+^ memory T cells associated with strong protection against SARS-CoV-2 infection, COVID19-like symptoms, and death.

## RESULTS

1. *Treatment of SARS-CoV-2-infected K18-hACE2 mice with CXCL-11 T-cell-attracting chemokine improves COVID-19-like symptoms and survivals:* We first determined whether treatment of SARS-CoV-2 infected K18-hACE2 mice with CXCL-9, CXCL-10, and CXCL-11 T-cell-attracting chemokines will improve COVID-19-like symptoms. Eight to nine-week-old K18-hACE2 transgenic mice (*n* = 20) were first infected intranasally with SARS-CoV-2 (1 x 10^4^ pfu/mouse of Washington USA-WA1/2020 variant) and were subsequently left untreated (control) or treated intranasally on days 3, 5, 7 and 9 post-infection either with CXCL-9, CXCL-10, or CXCL-11 chemokine (**Fig. 1A**). Chemokine-treated and untreated mice were then followed daily for the percentage of body weight and survival (i.e., COVID-19-like symptoms and death scoring). SARS-CoV-2-infected and CXCL-11-treated K18-hACE2 mice presented significant protection against weight loss (**Fig. 1B**) and death (**Fig. 1C**) compared to the SARS-CoV-2-infected untreated control K18-hACE2 mice (*P* < 0.05). However, SARS-CoV-2-infected and CXCL-9-treated K18-hACE2 mice did not show significant improvement in weight loss (**Fig. 1B**) and death (**Fig. 1C**) compared to the SARS-CoV-2-infected untreated control K18-hACE2 mice (*P* > 0.05). Similar to the CXCL-11-treated K18-hACE2 mice, the CXCL-10-treated K18-hACE2 mice showed better survival (100% survival vs. 75% survival, respectively, **Fig. 1C**). Altogether, these results indicate that among the three CXCL-9, CXCL-10, and CXCL-11 T-cell-attracting chemokines; treatment with the CXCL-11 chemokine significantly improved COVID-19-like symptoms and reduced deaths in SARS-CoV-2 infected K18-hACE2 mice.
2. *A multi-epitope/CXCL11 prime/pull Coronavirus vaccine protects against COVID19-like symptoms in HLA-DR*0101/HLA-A*0201/hACE2 triple transgenic mice following infection with SARS-CoV-2:* Since treatment with the CXCL-11, but not CXCL-9 and CXCL-10 chemokines, appeared to improve COVID-19-like symptoms and survivals of K18-hACE2 mice following infection with SARS-CoV-2, we next determined whether CXCL-11 treatment would also improve the protection induced by a multi-epitope coronavirus vaccine. We thus compared the protective efficacy of multi-epitope prime/pull Coronavirus vaccine candidates bearing multiple human CD4^+^ and CD8^+^ T cell epitopes. For this experiment, we: (1) designed and produced a multi-epitope Coronavirus vaccine that co-express recently identified 16 highly conserved human CD8^+^ T cell epitopes, 6 highly conserved human CD4^+^ T cell epitopes, and 8 highly conserved human B cell epitopes all expressed in tandem under improved CMV promoter (CAG) in a lung-tropic adeno-associated virus type 9 (AAV9) vector (designated as CoV-Vacc, **Fig. 2A**); and (2) generated a novel HLA-DR*0101/HLA-A*0201/hACE2 triple transgenic mice expressing human ACE2, human HLA class 1 (HLA-A*0201) and class 2 (HLA-DR*0101) following challenge with SARS-CoV-2. As illustrated in **Figs. 2B**, the prime/pull Coronavirus vaccination consists of (i) First priming of SARS-CoV-2-specific B cells, CD4^+^ T cells, and CD8^+^ T cells in HLA-DR*0101/HLA-A*0201/hACE2 triple transgenic mice using the engineered lung-tropic AAV9 multi-epitope Coronavirus vaccine co-expressing the recently identified immunodominant B, CD4^+,^ and CD8^+^ T cell epitopes (i.e. CoV-Vacc) and delivered intranasally followed by (ii) Pulling the “primed” B cells, CD4^+^ T cells, and CD8^+^ T cells into the lungs using mouse CXCL-9, CXCL-10, or CXCL-11 T-cell attracting chemokines delivered intranasally (nose drops). As illustrated in **Fig. 2B**, the HLA-DR*0101/HLA-A*0201/hACE2 triple transgenic mice (eight to nine-week-old, *n* = 35)) were intranasally immunized with 2×10^10^ VP per mouse of CoV-Vacc. The immunized mice were divided into 5 groups of 7 mice each and subsequently left untreated (CoV-Vacc. alone) or treated intranasally with 2.4 μg of CXCL-9 (*n* = 7), CXCL-10 (*n* = 7), or CXCL-11 (*n* = 7) on days 10, 12, 14, 22, 24, 26 post-immunization. The fifth control group (*n* = 7) was neither immunized with the CoV-Vacc. nor treated with the chemokines (mock). On day 28 post-immunization, mice were intranasally challenged with 1 x 10^4^ pfu of SARS-CoV-2 (USA-WA1/2020). Subsequently, all vaccinated groups received additional treatment with CXCL-9, CXCL-10, or CXCL-11 on days 30, 32, and 34 post-immunization. All animals were then monitored for up to day 14 post-infection (p.i.), for weight loss, virus replication in the lungs, and death. On day 14 p.i., mice were euthanized, and lungs were collected for lung inflammation using H & E staining. We observed significant protection against weight loss in the group of mice that received the multi-epitope//CXCL11 prime/pull CoV-Vacc compared to the multi-epitope//CXCL9 prime/pull CoV-Vacc. group, the multi-epitope//CXCL10 prime/pull CoV-Vacc group, as well as the Mock control group (*P* < 0.005, **Fig. 3A**). Between days 5 and 9 post-challenge with SARS-CoV-2, the multi-epitope//CXCL11 prime/pull CoV-Vacc group lost only around 2% of their initial weight, whereas the remaining groups showed a significant loss in their initial weight that varied between 4% to 10% (**Fig. 3A**). Accordingly, a significant reduction in virus replication was recorded in the lungs of multi-epitope//CXCL11 prime/pull CoV-Vacc group of mice compared to other groups (**Fig. 3C**). Significantly lower viral RNA copy numbers were detected on both days 4 and 8 post-infection in the lungs of the multi-epitope//CXCL11 prime/pull CoV-Vacc group compared to other groups (**Fig. 3C**). Moreover, from day 0 to day 14 post-challenge with SARS-CoV-2, the multi-epitope//CXCL11 prime/pull CoV-Vacc group of mice showed 100% survival whereas the remaining groups depicted only 60% to 80% survival (**Fig. 3B**). The lowest virus replication in the lungs of multi-epitope//CXCL11 prime/pull CoV-Vacc. group of mice was associated with less inflammatory cells infiltrating the lungs (**Fig. 3D**). Less pulmonary pathological changes characterized by (i) open alveolar air spaces, (ii) less inflammation, and (iii) less residual cellular debris in air spaces with less alveolar damage, were observed in the H & E sections from mice immunized with Pan-CoV-Vacc and treated with CXCL-11 (**Fig. 3D**). Altogether, these results demonstrate that the multi-epitope//CXCL11 prime/pull Coronavirus vaccine protected against COVID19-like symptoms and reduced virus replication and inflammation in the lungs, and prevented deaths in HLA-DR*0101/HLA-A*0201/hACE2 triple transgenic mice following infection with SARS-CoV-2.
3. *Bolstering the frequencies of lung-resident memory CD4^+^ and CD8^+^ T_EM_, T_CM,_ and T_RM_ cells through the multi-epitope/CXCL11 prime/pull Coronavirus vaccine protected HLA-DR*0101/HLA-A*0201/hACE2 triple transgenic mice following SARS-CoV-2 infection:* Since the multi-epitope//CXCL11 prime/pull Coronavirus vaccine appeared to prevent weight loss, to reduce virus replication and inflammation in the lungs and to prevent deaths in HLA-DR*0101/HLA-A*0201/hACE2 triple transgenic mice following infection with SARS-CoV-2, we next determined whether such protection would be associated with increased frequencies of lung-resident memory CD4^+^ and CD8^+^ T cells. Next, we compared the effect of immunization with the three multi-epitope prime/pull Coronavirus vaccine candidates based on CXCL-9, CXCL-10, and CXCL-11 T-cell-attracting chemokines on the frequencies of three major lung-resident memory CD4^+^ and CD8^+^ T cell subsets (i.e. effector memory (T_EM_), resident memory (T_RM_), and central memory (T_CM_)) in the lungs of HLA-DR*0101/HLA-A*0201/hACE2 triple transgenic mice, before (**Fig. 4**) and after (**Fig. 5**) challenge with SARS-CoV-2 (**Fig. 2B**). As controls, the frequencies of three major lung-resident memory CD4^+^ and CD8^+^ T cell subsets were compared in HLA-DR*0101/HLA-A*0201/hACE2 triple transgenic mice that received the same multi-epitope CoV-Vacc bearing human CD4^+^ and CD8^+^ T cell epitopes without chemokine treatment (*CoV Vacc alone*) as well as in mock-vaccinated mice (*Mock*). For this experiment, eight to nine-week-old HLA-DR*0101/HLA-A*0201/hACE2 triple transgenic mice (*n* = 35) were first immunized with the multi-epitope Coronavirus vaccine (CoV-Vacc) delivered intranasally at 2 x 10^10^ VP per mouse. Vaccinated HLA-DR*0101/HLA-A*0201/hACE2 triple transgenic were subsequently left untreated (control) or treated intranasally on days 10, 12, and 14 post-vaccination and on days 22, 24, and 26 post-vaccination with the CXCL-9, CXCL-10, or CXCL-11 T-cell-attracting chemokine as described in *Materials* and *Methods*. Vaccinated/chemokine-treated and Vaccinated/untreated mice were euthanized on day 27 post-vaccine and the frequencies of the three major lung-resident memory CD4^+^ and CD8^+^ T cells expressing CXCR3, CD103, CD62L, and CD44 among total lung cells (i.e. effector memory (T_EM_), resident memory (T_RM_), and central memory (T_CM_)) were determined by FACS, as described in *Materials* and *Methods*. We found a significant increase in the frequency of total CD4^+^ T cells among immunized mice compared to mock mice (**Fig. 4A**). Similarly, we found a significant increase in CD103^+^CD4^+^ T cells, CD44^+^CD62L^-^ CD4^+^ T cells, and CD44^+^CD62L^+^CD4^+^ T cells in mice immunized with the CoV-Vacc. However, CXCR3^+^CD4^+^ T cells did not show significant variation for mice immunized with CoV Vacc + CXCL-9, CoV Vacc + CXCL-10, or CoV Vacc + CXCL-11, compared to mice immunized only with Vacc or mock group of mice. In (**Fig. 5A**), a higher magnitude in the frequency of total CD8^+^ T cells in lung immune cells was shown to be observed in the immunized mice. Further, the mice immunized with CoV Vacc showed a significant increase in CXCR3^+^CD8^+^ T cells, CD44^+^CD62L^-^CD8^+^ T cells, and CD44^+^CD62L^+^CD8^+^ T cells when compared to the mock group. However, CD103^+^CD8^+^ T cells showed a decrease in the CoV Vacc + CXCL-11 group compared to the immunized and mock groups. Altogether, these results demonstrate that the protection induced by the multi-epitope//CXCL11 prime/pull Coronavirus vaccine in SARS-CoV-2 infected HLA-DR*0101/HLA-A*0201/hACE2 triple transgenic mice is associated with high frequencies of lung-resident memory CD4^+^ and CD8^+^ T_EM_, T_CM,_ and T_RM_ cells.
4. *Bolstering the function of virus-specific lung-resident memory CD4^+^ and CD8^+^ T cells through the multi-epitope/CXCL11 prime/pull Coronavirus vaccine protected HLA-DR*0101/HLA-A*0201/hACE2 triple transgenic mice following SARS-CoV-2 infection::* Since the protection induced by the multi-epitope//CXCL11 prime/pull Coronavirus vaccine in SARS-CoV-2 infected HLA-DR*0101/HLA-A*0201/hACE2 triple transgenic mice is associated with high frequencies of lung-resident memory CD4^+^ and CD8^+^ T_EM_, T_CM,_ and T_RM_ cells, we next determined whether such protection would be associated with increased frequencies of lung-resident memory CD4^+^ and CD8^+^ T cells.

We compared the effect of immunization with the three multi-epitope prime/pull Coronavirus vaccine candidates based on CXCL-9, CXCL-10, and CXCL-11 T-cell-attracting chemokines on the function and specificity of pulled CD4^+^ and CD8^+^ T cells in the lungs of HLA-DR*0101/HLA-A*0201/hACE2 triple transgenic mice, before (**Fig. 6**) challenge with SARS-CoV-2 (**Fig. 2B**). As controls, HLA-DR*0101/HLA-A*0201/hACE2 triple transgenic mice that received the same multi-epitope CoV-Vacc bearing human CD4^+^ and CD8^+^ T cell epitopes without chemokine treatment (*CoV Vacc alone*) as well as in mock-vaccinated mice (*Mock*).

**Figure 1.**
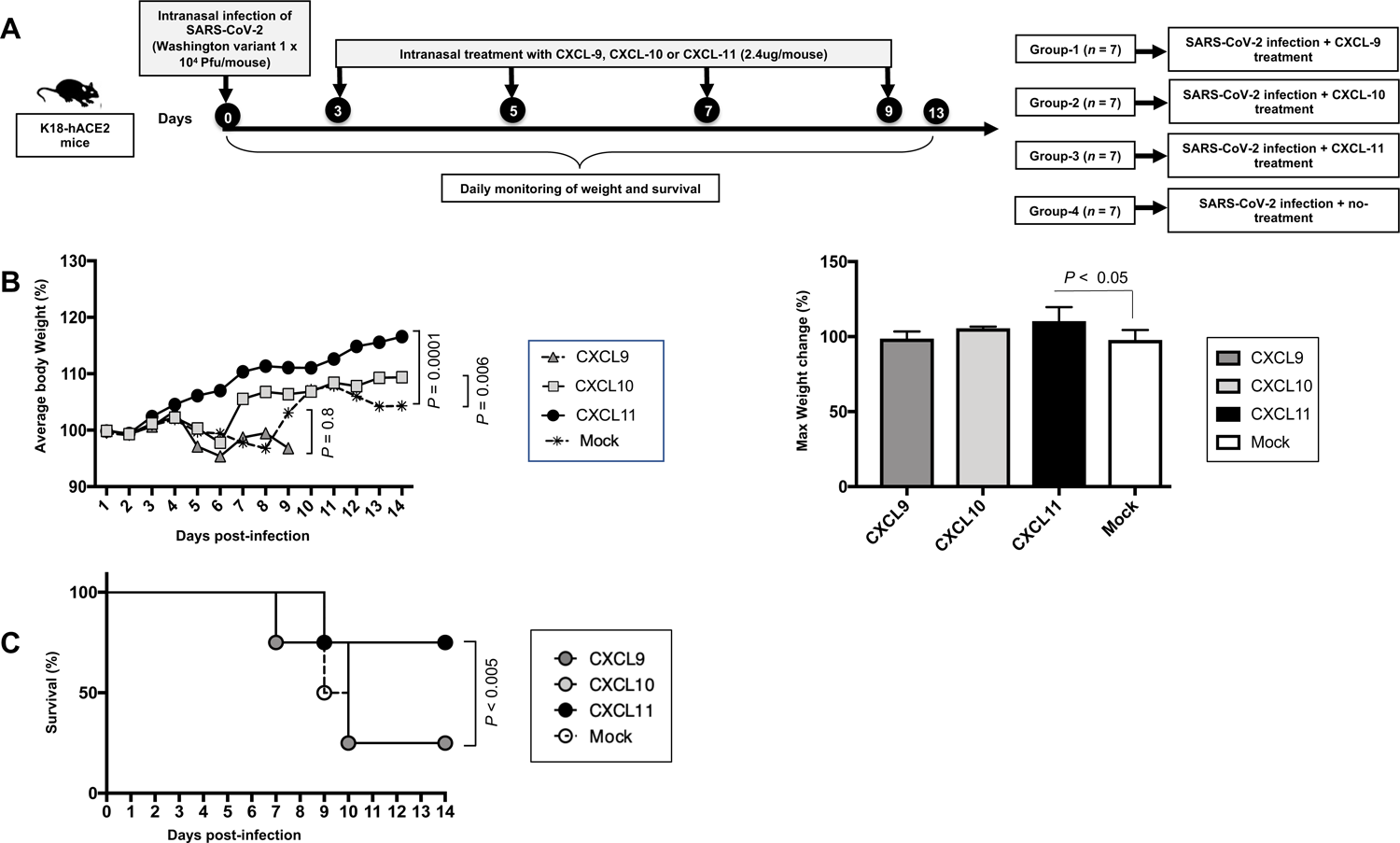
The effect of treatment with CXCL-9, CXCL-10, and CXCL-11 chemokines on COVID-19-like symptoms detected from K18-hACE2 single transgenic mice infected with SARS-CoV-2. (**A**) Experimental plan to study the effect of treatment with CXCL-9, CXCL-10, and CXCL-11 chemokines on COVID-19-like symptoms detected from K18-hACE2 single transgenic mice following infection with SARS-CoV-2 (Washington USA-WA1/2020 variant). 8-9 week-old male and female K18-hACE2 mice (*n* = 20) were infected intranasally with 1 x 10^4^ pfu of the SARS-CoV-2-USA-WA1/2020 variant. Mice were subsequently treated intranasally with CXCL-9 (*n* = 4), CXCL-10 (*n* = 4), and CXCL-11 (*n* = 4) on days 3, 5, 7, and 9 post-infection (p.i). As a control mice (*n* = 4) were left untreated. (**B**) Graph shows average body weight change p.i. normalized to the body weight on the day of infection (*right panel*). The maximum percent body weight change on day 7 p.i. is shown in the *left panel.* (**C**) Shows the percentage survival detected in CXCL-9, CXCL-10, and CXCL-11 treated vs. untreated mice up to day 13 p.i.

**Figure 2.**
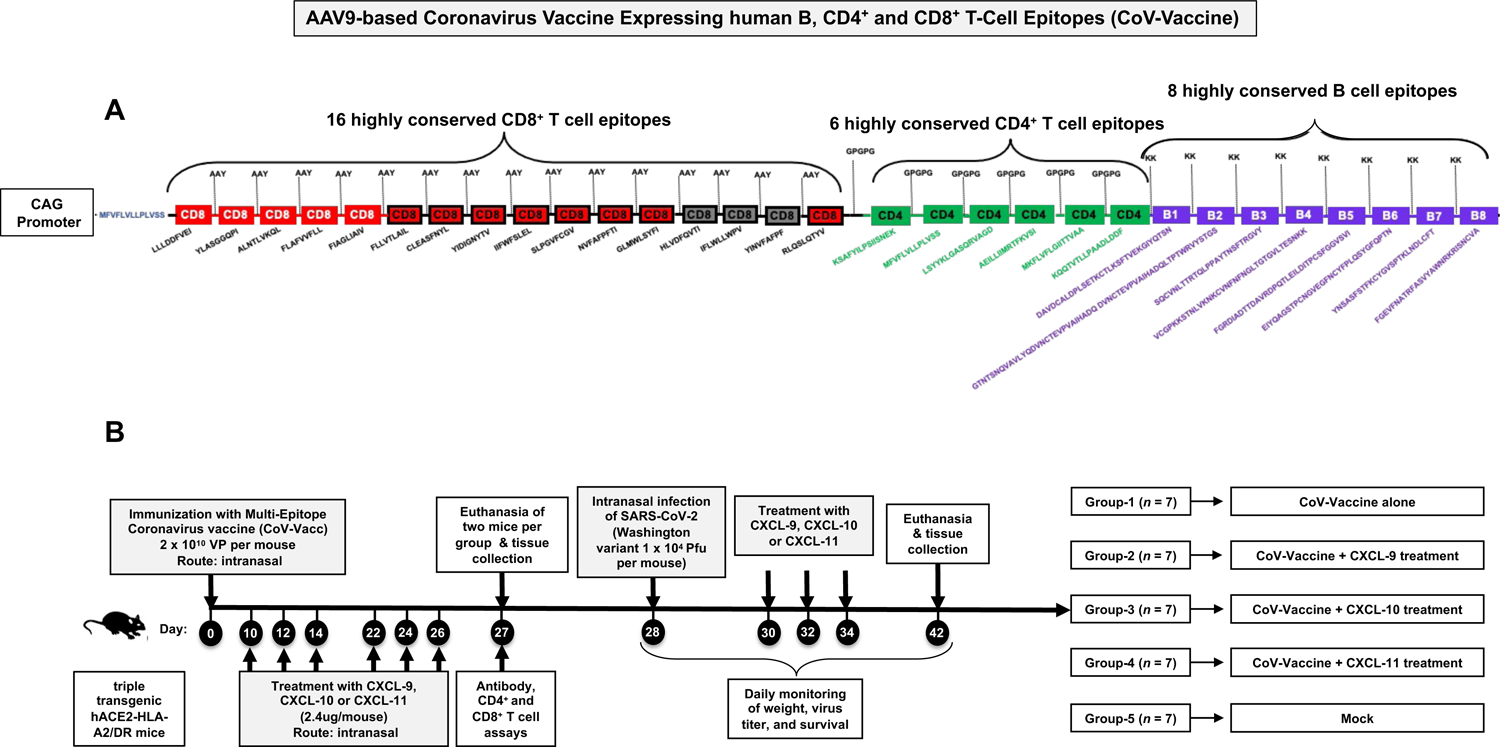
The effect of treatment with CXCL-9, CXCL-10, and CXCL-11chemokines on disease outcome detected from HLA-DR*0101/HLA-A*0201/hACE2 triple transgenic mice immunized with SARS-CoV-2 vaccine. (**A**) Figure showing a prototype of the multi-epitope Coronavirus vaccine consisting of highly conserved and immunogenic 16 CD8^+^ T cell epitopes, 6 CD4^+^ T cell epitopes, and 8 B cell epitopes. (**B**) Experimental plan to study the effect of treatment with CXCL-9, CXCL-10, and CXCL-11 chemokines on COVID-19-like symptoms detected from HLA-DR*0101/HLA-A*0201/hACE2 triple transgenic mice immunized with multi-epitope Coronavirus vaccine (CoV-Vacc). 8-9 week-old male and female HLA-DR*0101/HLA-A*0201/hACE2 triple transgenic mice were intranasally immunized with CoV-Vacc; 2×10^10^ VP per mice on day 0 (*n* = 28). As a control mice (*n* = 7) were left unimmunized. The immunized mice were subsequently treated intranasally with 2.4 μg of; CXCL-9 (*n* = 7), CXCL-10 (*n* = 7), and CXCL-11 (*n* = 7) on days 10,12,14,22,24,26 post-immunization. On day 27 two mice per group were euthanized, and lung tissues were collected. The immune cell response was evaluated by flow cytometry. The remaining mice (*n* = 25) were intranasally infected with 1×10^4^ pfu of SARS-CoV-2 (USA-WA1/2020) on day 28 post-immunization. Three mice groups were subsequently treated intranasally with CXCL-9 (*n* = 5), CXCL-10 (*n* = 5), and CXCL-11 (*n* = 5) on days 30, 32, and 34 post-immunization. disease monitoring, weighing, and survival were monitored in the mice up to day 14 p.i.

**Figure 3.**
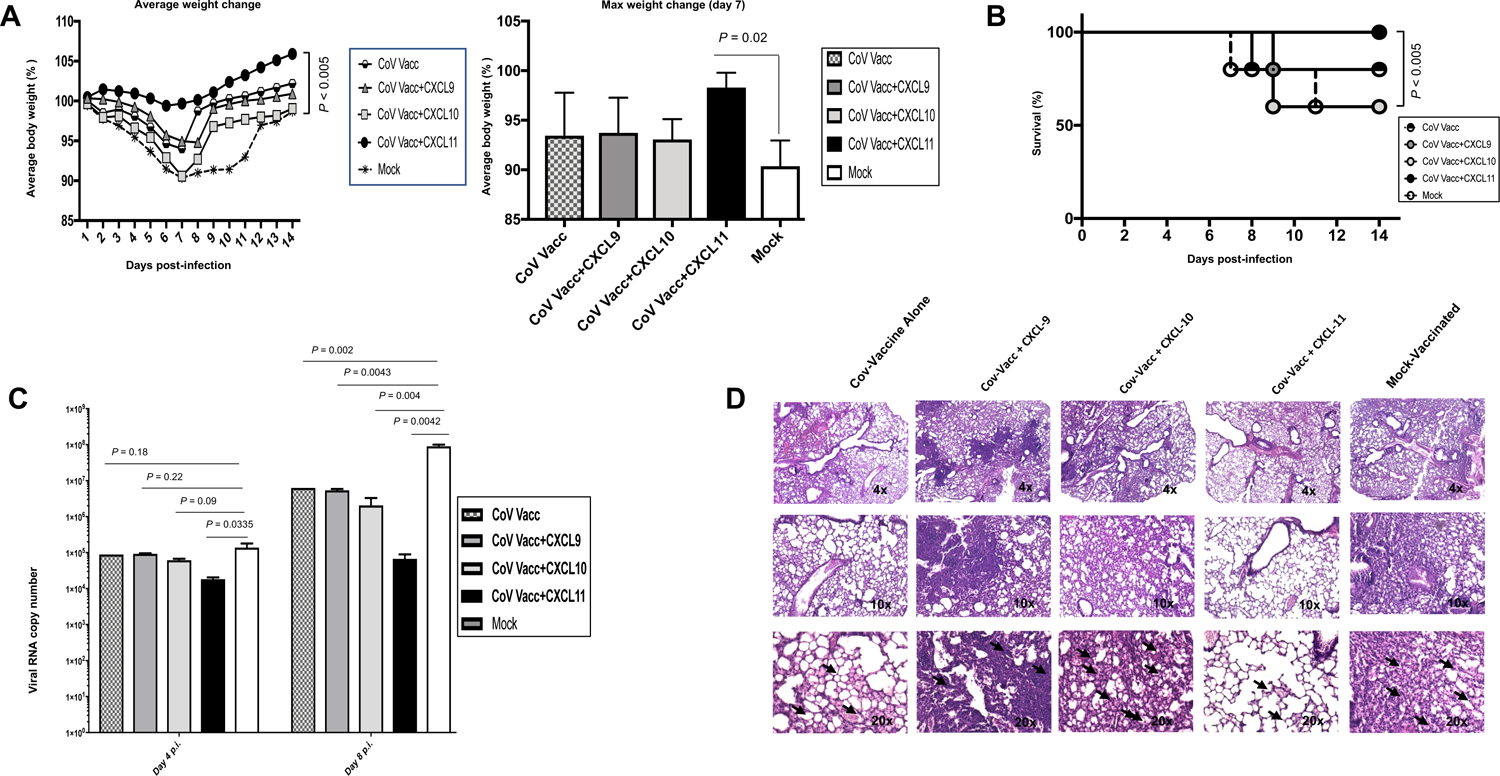
The effect of treatment with CXCL-9, CXCL-10, and CXCL-11 chemokines on COVID-19-like symptoms detected from HLA-DR*0101/HLA-A*0201/hACE2 triple transgenic mice immunized with multi-epitope Coronavirus vaccine and challenged with SARS-CoV-2-USA-WA1/2020 variant. (**A**) Data showing average percent weight change each day p.i. normalized to the body weight on the day of infection is shown in the *right panel*. The bar graph (*left panel*) shows the percent weight change at day 7 p.i. Bars represent means ± SEM. (**B**) Shows the percentage survival detected in mice groups of, CoV-Vacc, CoV-Vacc + CXCL-9, CoV-Vacc + CXCL-10, CoV-Vacc + CXCL-11, and mock vaccinated group up to 14 days p.i. (**C**) Viral titration data showing viral RNA copy number in the lungs for each group at days 4 and 8 p.i. (**D**) Representative H & E staining images of the lungs at day 14 p.i. of SARS-CoV-2 infected mice treated with different chemokines at 4x, 10x, and 40x magnifications.

**Figure 4.**
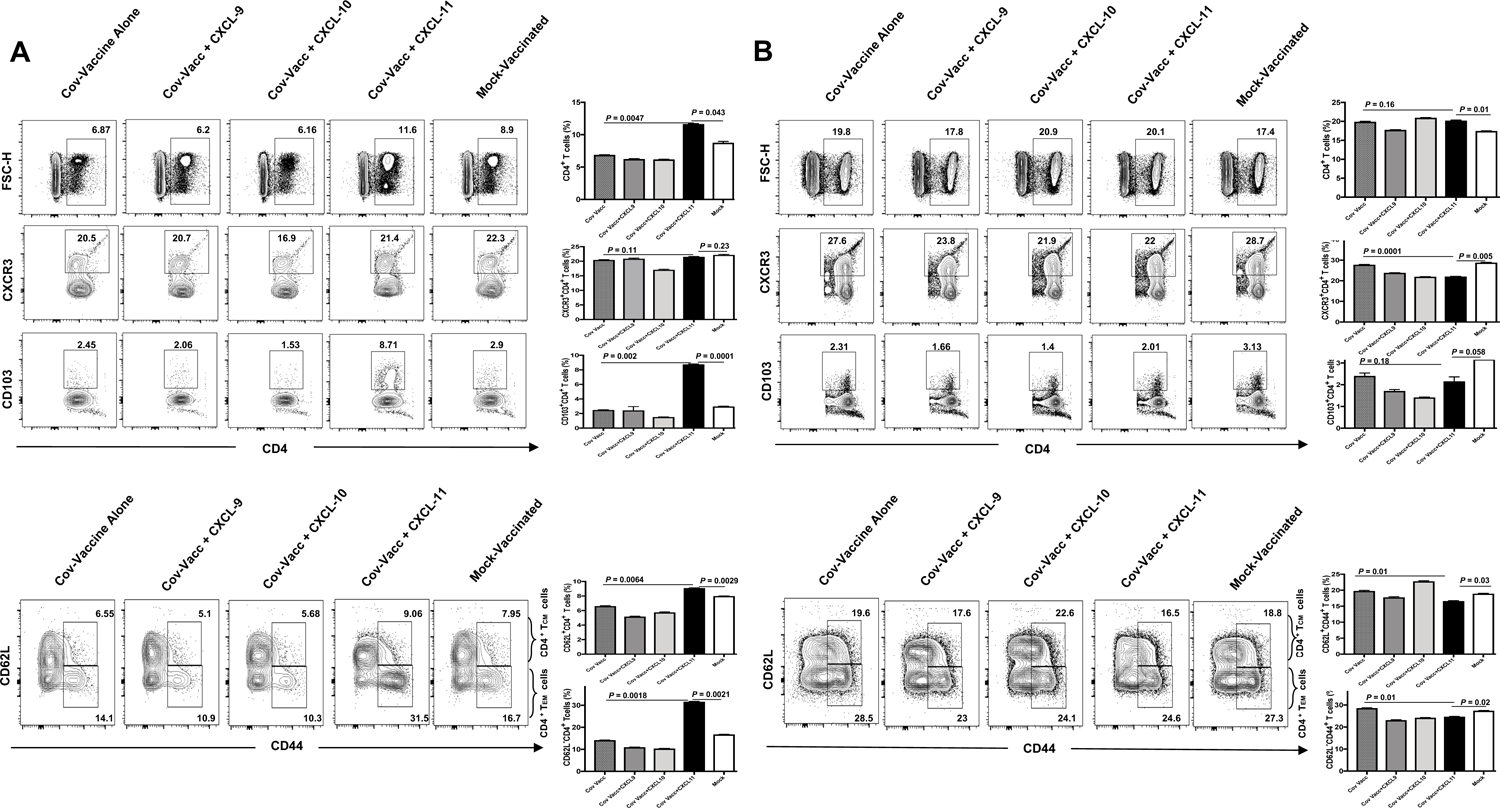
The effect of treatment with CXCL-9, CXCL-10, and CXCL-11chemokines on CD4^+^ T cells in the lung and spleen of immunized HLA-DR*0101/HLA-A*0201/hACE2 triple transgenic mice. (**A**) The left *panel* shows FACS plots for CD4^+^ T cells in the lungs of mice immunized with the multi-epitope Coronavirus vaccine (*n* = 7) and treated with CXCL-9, CXCL-10, and CXCL-11. Graphs depict the differences in response to various treatments on the percentage of CD4^+^ T cells present in the lungs of mice shown in the *right panel*. Bars represent the means ± SEM. Student’s t-test was used to analyze the data. (**B**) The left *panel* represents FACS plots for CD4^+^ T cells in the spleen of mice immunized with the multi-epitope Coronavirus vaccine (*n* = 7) and treated with CXCL-9, CXCL-10, and CXCL-11. Graphs on the *right panel* show the difference in response to different chemokine treatments on the percentage of CD4^+^ T cells in the spleen of mice. Bars represent means ± SEM.

**Figure 5.**
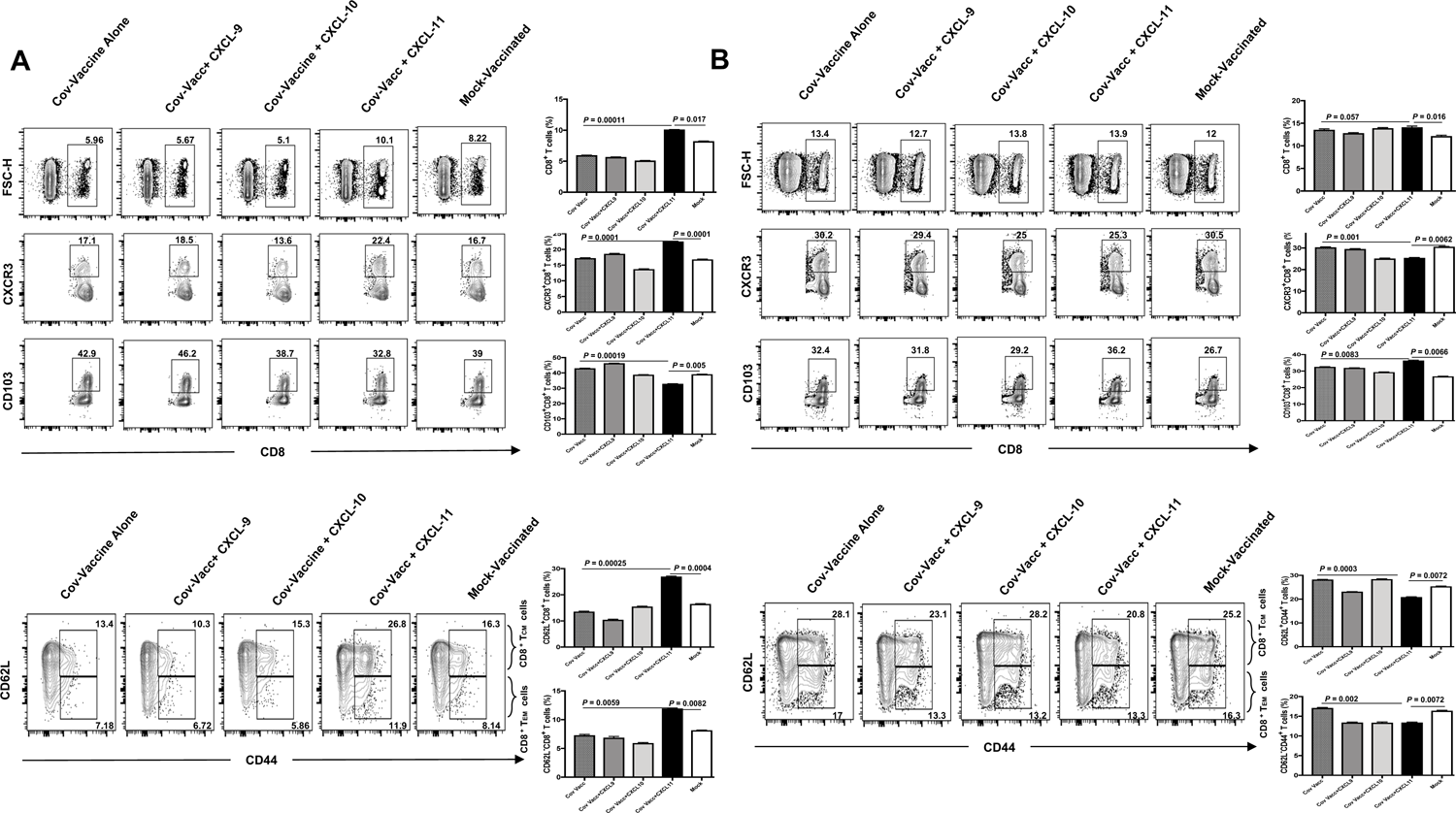
The effect of treatment with CXCL-9, CXCL-10, and CXCL-11chemokines on CD8^+^ T cells in the lung and spleen of immunized hACE2-HLA-A2/DR triple transgenic mice. (**A**) The left *panel* shows FACS plots for CD8^+^ T cells in the lungs of mice immunized with the multi-epitope Coronavirus vaccine (*n* = 7) and treated with CXCL-9, CXCL-10, and CXCL-11. Graphs depict the differences in response to various treatments on the percentage of CD8^+^ T cells present in the lungs of mice shown in the *right panel*. Bars represent the means ± SEM. Student’s t-test was used to analyze the data. (**B**) The left *panel* represents FACS plots for CD8^+^ T cells in the spleen of mice immunized with the multi-epitope Coronavirus vaccine (*n* = 7) and treated with CXCL-9, CXCL-10, and CXCL-11. Graphs on the *right panel* show the difference in response to different chemokine treatments on the percentage of CD4^+^ T cells in the spleen of mice. Bars represent means ± SEM.

**Figure 6.**
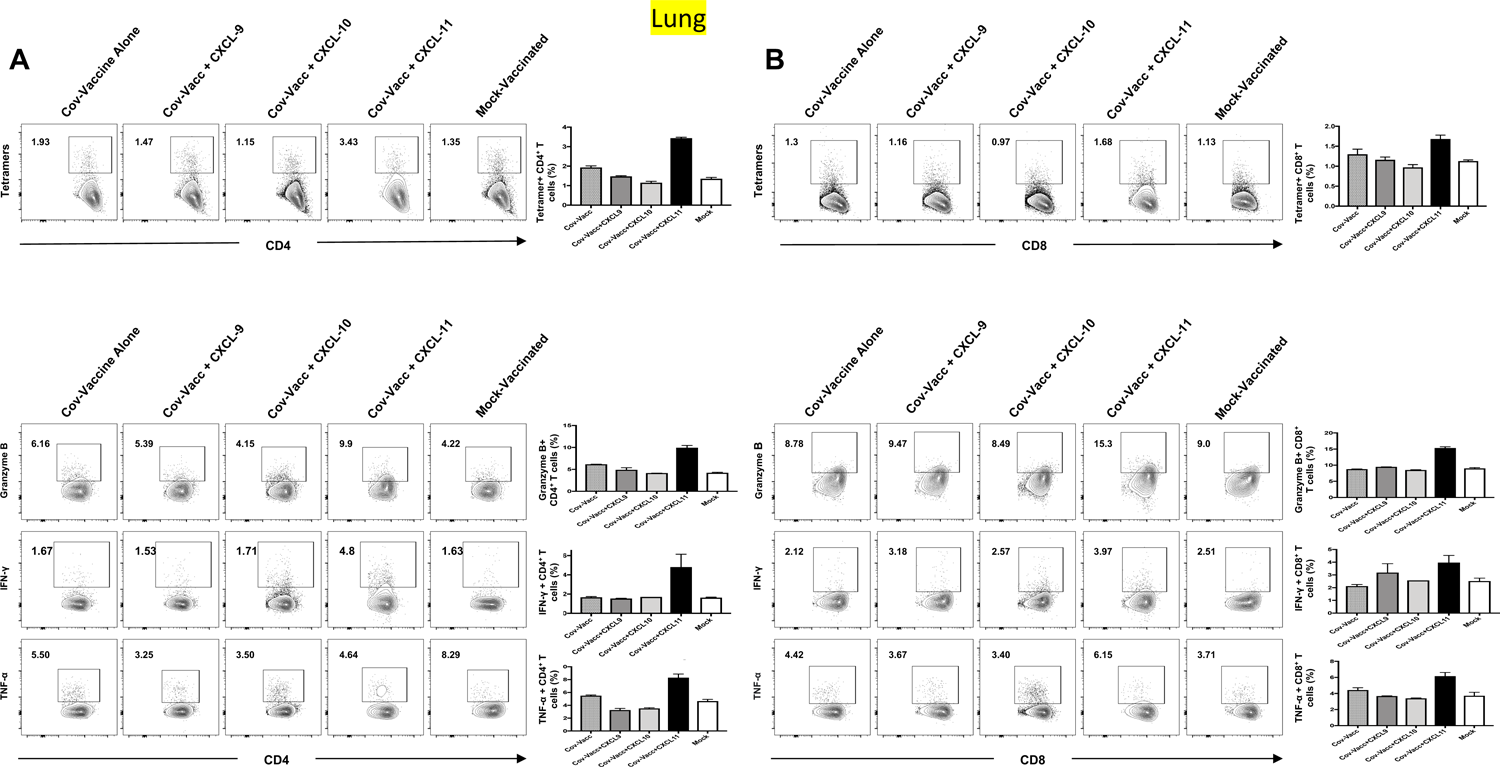
The effect of treatment with CXCL-9, CXCL-10, and CXCL-11chemokines on CD4^+^ T cells and CD8^+^ T cells in the lung of immunized hACE2-HLA-A2/DR triple transgenic mice. (**A**) The left *panel* shows FACS plots for CD4^+^ T cells in the lungs of mice immunized with the multi-epitope Coronavirus vaccine (*n* = 5) and treated with CXCL-9, CXCL-10, and CXCL-11. Graphs depict the differences in response to various treatments on the percentage of CD4^+^ T cells present in the lungs of mice shown in the *right panel*. Bars represent the means ± SEM. Student’s t-test was used to analyze the data. (**B**) The left *panel* represents FACS plots for CD8^+^ T cells in the lungs of mice immunized with the multi-epitope Coronavirus vaccine (*n* = 5) and treated with CXCL-9, CXCL-10, and CXCL-11. Graphs on the *right panel* show the difference in response to different chemokine treatments on the percentage of CD8^+^ T cells in the lungs of mice. Bars represent means ± SEM.

For this experiment, eight to nine-week-old HLA-DR*0101/HLA-A*0201/hACE2 triple transgenic mice (*n* = 35) were first immunized with the multi-epitope Coronavirus vaccine (CoV-Vacc) delivered intranasally at 2 x 10^10^ VP per mouse. Vaccinated HLA-DR*0101/HLA-A*0201/hACE2 triple transgenic were subsequently left untreated (control) or treated intranasally on days 10, 12, and 14 post-vaccination and on days 22, 24, and 26 post-vaccination with the CXCL-9, CXCL-10, or CXCL-11 T-cell-attracting chemokine as described in *Materials* and *Methods*. Vaccinated/chemokine-treated and Vaccinated/untreated mice were euthanized on day 27 post-vaccine immune cells were harvested for the frequency and the function of major lung-memory CD4^+^ and CD8^+^ T cells determined by FACS, as described in *Materials* and *Methods*.

We found a significant increase in the percentage of tetramer^+^CD4^+^ T cells among immunized and CXCL11-treated mice compared to all other mice groups (**Fig. 6A**). Similarly, we found a significant increase in Granzyme B^+^CD4^+^ T cells, IFN-γ^+^CD4^+^ T cells, and TNF-α^+^CD4^+^ T cells in mice immunized with the CoV-Vacc and treated with the chemokine CXCL11. In (**Fig. 6B**), a slight increase in magnitude in the percentage of tetramer^+^CD8^+^ T cells in the lungs of immunized and CXCL11-treated mice compared to all other mice groups. A significant increase in Granzyme B^+^CD8^+^ T cells and TNF-α^+^CD8^+^ T cells was shown in the lungs of the CoV Vacc + CXCL-11 group compared to the immunized and mock groups. While IFN-γ^+^CD8^+^ T cells didn’t show a significant increase among all the groups

Altogether, these results demonstrate that the protection induced by the multi-epitope//CXCL11 prime/pull Coronavirus vaccine in SARS-CoV-2 infected HLA-DR*0101/HLA-A*0201/hACE2 triple transgenic mice is associated with high frequencies of specific of CD4^+^ and CD8^+^ T cells in the lungs. As well as high expression of Granzyme B, IFN-γ, and TNF-α of those cells.

## DISCUSSION

For more than 3 years, humanity has been confronting the COVID-19 pandemic caused by the new SARS-CoV-2 infection (20). While current parenteral COVID-19 parenteral vaccines have lessened immediate threats by COVID-19, they induced low frequencies of memory SARS-CoV-2-specific CD4^+^ and CD8^+^ T cells in the lungs. Unlike mucosal vaccines, parenteral vaccines are often less effective in inducing mucosal T-cell immunity, such as in the lungs, the site of virus replication (21). Thus, the low frequencies of memory SARS-CoV-2-specific CD4^+^ and CD8^+^ T cells in the lungs are associated with severe cases of acute and long COVID-19, even in some vaccinated patients (22). Moreover, because most mutations and deletions that produced the 20 known variants of concern (VOCs) are concentrated within the sequence of the Spike protein, there is a risk that the current Spike-based COVID-19 sub-unit vaccines will fail to protect against future VOCs despite inducing strong neutralizing antibodies against the original virus strain (55). This emphasizes the need for second-generation Coronavirus vaccines that: (1) Target antigens (Ags) other than the highly variable Spike protein; (2) Incorporate both B- and T-cell epitopes from Spike and non-Spike Ags that are highly conserved in all 20 VOCs and that will induce strong humoral and cell-mediated immune responses; and (3) Can boost the frequencies of functional SARS-CoV-2-specific CD4^+^ and CD8^+^ T cells in the lungs.

In the present study, we describe a novel prime/pull Coronavirus mucosal vaccine strategy that consists of intranasal delivery of a multi-epitope Coronavirus vaccine that incorporates genome-wide highly conserved SARS-CoV-2 B, CD4^+^ CD8^+^ cell epitopes (*prime*), and pulling primed T cells using the CXCL-11 T-cell-attracting chemokine (*pull*). We demonstrated that such a multi-epitope//CXCL11 prime/pull Coronavirus mucosal vaccine: (1) Increased the frequencies of functional CD4^+^ and CD8^+^ T_EM_, T_CM,_ and T_RM_ cells in the lungs of HLA-DR*0101/HLA-A*0201/hACE2 triple transgenic mice; (2) protected against COVID19-like symptoms, reduced virus replication in the lungs, and prevented deaths following intranasal inoculation with SARS-CoV-2. We discussed the new prime/pull vaccine strategy and the importance of boosting the lung-resident SARS-CoV-specific CD4^+^ and CD8^+^ memory T cell subsets for better protection against SARS-CoV-2 infection, COVID-19-like symptoms, and death.

The current parenteral subunit COVID vaccines that rely on neutralizing Spike-specific antibodies have prevented SARS-CoV-2-related hospital admission and deaths. However, many Spike-alone-based vaccines induced low frequencies of memory SARS-CoV-2-specific CD4^+^ and CD8^+^ T cells in the lungs associated with severe cases of acute and long COVID-19, even in some boosted patients (23). To address this issue, we demonstrated a novel multi-epitope//CXCL11 prime/pull Coronavirus mucosal vaccine strategy boosted the frequencies of functional CD4^+^ and CD8^+^ T_EM_, T_CM,_ and T_RM_ cells in the lungs of HLA-DR*0101/HLA-A*0201/hACE2 triple transgenic mice associated with strong protection against SARS-CoV-2 infection, COVID-19-like symptoms, and death. In contrast, such protection did not correlate with blocking/neutralizing antibodies. This highlights the importance of lung-resident CD4^+^ and CD8^+^ memory T cells in protection against SARS-CoV-2 infection, COVID-19-like symptoms, and death (24). Moreover, T cell-based vaccines are critical in SARS-CoV-2 as they can provide long-lasting memory immunity that could potentially protect against repetitive infection with multiple variants. CD4^+^ and CD8^+^ memory T cells can recognize and eliminate infected cells and provide memory immunity in case of re-exposure (25, 26). In addition, unlike antibody-based vaccines, T-cell-based vaccines have the potential to protect against new and mutated strains of the virus (27).

Chemokines play a complex and significant role in the immune response to viral infections, including SARS-CoV-2, by stimulating the production of antiviral proteins, such as interferons (28). Studies have shown that certain chemokines, such as CXCL-10, can stimulate the production of interferons in response to SARS-CoV-2 infection (52). Explicitly, the CXCL 10/CXCR3 pathways are imperative in guarding against viral pathogens through the means of T-cell immunity (29). While they can help to recruit immune cells to the site of infection and stimulate the production of antiviral proteins, excessive or dysregulated production of certain chemokines may also contribute to the development of severe COVID-19 (30). Hyperinflammation and tissue impairment could reside as a result of the vast attraction of cells towards the area of infection; the cytotoxic nature of chemokines has shown to have both advantageous and disadvantageous responses to viral infection (31). However, the explicit roles of these chemokines in SARS-CoV-2 immunity remain to be fully explored. Therefore, chemokines and their receptors are important targets for the development of therapeutics to treat different infectious diseases including SARS-CoV-2. Coperchini et. al. has identified 53 immune responses that involve cytokine activation; these chemokines are affiliated with immune cell trafficking, respiratory infection regulation, and the chemoattraction of T cells (32). The prime-pull approach has been used in the development of many successful vaccines, including those against influenza, HPV, and hepatitis B (33). The objective of our study was to develop an immunization strategy that protects against COVID-19-like symptoms and achieves a robust and long-lasting immune response.

The present study demonstrates a differential role for the CXCL-9, CXCL-10, and CXCL-11 chemokines in the mobilization of memory CD4^+^ and CD8^+^ T cells into the lungs and showed association of high frequencies of memory CD4^+^ and CD8^+^ T cells with protection against SARS-CoV-2 infection, COVID19-like symptoms, and death. We compared differential effects from treatment with CXCL-9, CXCL-10, and CXCL-11 chemokines on COVID-19-like symptoms detected from K18-hACE2 single transgenic mice following infection with SARS-CoV-2 (Washington USA-WA1/2020 variant). Mice that were treated with CXCL-11 had slightly better, but not significant protection against weight loss compared to the control group. However, mice that were treated with CXCL-9 and CXCL-10 did not depict any significant differences compared to the control group. The group of mice treated with CXCL-11 and CXCL-10 showed better survival with a protection of 75%, and mice treated with CXCL-9 showed a 25% survival compared to control mice by day 13 p.i. These results demonstrate that the addition of chemokine treatment could greatly enhance protective immunity against SARS-CoV-2-like symptoms. Therefore, it suggests that at least one of the above-listed chemokines (CXCL-11) might be a potential target for the treatment of COVID-19-like symptoms.

Following acute infection, three major SARS-CoV-2-specific memory CD4^+^ and CD8^+^ T cell sub-populations, T_CM_, T_EM,_ and T_RM_, develop, infiltrate and sequester in the infected lungs in response to a high level of T cell attracting CXCL9, CXCL10 and CXCL11 chemokines and other factors. Memory CD4^+^ and CD8^+^ T cell infiltrates were reported in the infected lungs of COVID-19 human cadavers. Functional CD4^+^ and CD8^+^ T cells in these infiltrates likely help decrease viral replication in the lungs, the brain, and likely other compartments. However, the apparent low level of T cell-attracting CXCL9, CXCL10, and CXCL11 chemokines in infected lungs may not be sufficient enough to assure and the sequestration of CD4^+^ and CD8^+^ T_RM_ cells or to guide homing CD4^+^ and CD8^+^ T_EM_ and T_CM_ cells from the circulation into infected lungs. In this study, we found that local delivery of the T cell-attracting CXCL11 chemokine, but not of CXCL9 or CXCL10 chemokines, dramatically increased both the number and function of SARS-CoV-2-specific CD4^+^ and CD8^+^ T_EM_ and T_RM_ cell sub-populations in infected lungs, improving protection against virus replication, COVID19-like symptoms, and death in SARS-CoV-2-infected HLA-DR*0101/HLA-A*0201/hACE2 triple transgenic mice. These results suggest that: (*i*) the number and/or the function of CD4^+^ and CD8^+^ T cells specific for “ASYMP” epitopes is suppressed in infected human lungs; (*ii*) local delivery of the T cell-attracting CXCL11 chemokine will expand the repertoire and function of SARS-CoV-2-specific CD8^+^ T cells and reduce the likelihood of viral replication in the lungs; and (*iii*) a prime/pull vaccine strategy that increases the number and/or function of CD4^+^ and CD8^+^ T cells in infected lungs over a certain threshold would likely lead to protection in humans. The success of this innovative prime/pull vaccine is likely due to the expansion and survival characteristics of CD4^+^ and CD8^+^ T cell precursors as well as the increase in the number and function of other immune cells in lungs, including APCs, following CXCL11 treatment (34). This contributes not only to homing of CD4^+^ and CD8^+^ T_EM_ cell sub-populations in infected lungs but also to the increased expansion and survival of local tissue-resident CD4^+^ and CD8^+^ T_RM_ cells that already exist within the lungs.

SARS-CoV-specific CD4^+^ and CD8^+^ T cells that reside in the lungs appeared to play a critical role in aborting virus replication (35). In designing a human multi-epitope Coronavirus vaccine, it is critical to identify the protective Coronavirus epitopes derived from the 26^+^ ORFs of the SARS-COV-2 genome, which are exclusively recognized by the human CD4^+^ and CD8^+^ T cells from “naturally” protected ASYMP individuals who, despite being infected they never develop any severe COVID-19 disease (36). We previously implemented several *in silico, ex vivo*, *in vitro*, and *in vivo* phenotypic and functional immunological methods to efficiently generate a genome-wide map of the responsiveness of SARS-COV-2-specific CD4^+^ and CD8^+^ T cells in ASYMP individuals. This leads to the identification of several previously unknown protective human “ASYMP” CD4^+^ and CD8^+^ T cell epitopes. These “ASYMP” epitopes are associated with frequent, robust, and polyfunctional CD4^+^ and CD8^+^ T_EM_ cell responses. The present study extends previous findings by demonstrating that immunization of a novel “humanized” susceptible HLA-DR*0101/HLA-A*0201/hACE2 triple transgenic mouse model with “ASYMP” epitopes induced a strong CD4^+^ and CD8^+^ T cell-dependent protective immunity against COVID-19 like symptoms. Furthermore, we report a novel prime-pull vaccine strategy, based on priming CD4^+^ and CD8^+^ T cells with multiple “ASYMP” epitopes followed by treatment with the CXCL11 T cell-attracting chemokine. This approach significantly boosts the number of functional anti-viral CD4^+^ and CD8^+^ T_EM_ and T_RM_ cells in the lungs of SARS-CoV-2 infected humanized” HLA-DR*0101/HLA-A*0201/hACE2 triple transgenic mice and improves protection against COVID-like symptoms following SARS-CoV-2 infection. These findings have profound implications for the development of T-cell-based prophylactic COVID vaccine strategies to protect against COVID-19 infections and diseases in humans.

The development of multi-B- and T-cell epitope-based vaccines in the era of omics is taking advantage of new technologies to tackle other Coronavirus infections and diseases for which vaccine development has been unsuccessful (37). Cutting-edge technologies and screening strategies have recently been developed for many genomic sequence information for the state-of-the-art rational multi-B- and T-cell epitope-based vaccine design (38). However, low frequencies of SARS-CoV-2-specific CD4^+^ and CD8^+^T cells have hampered genome-wide identification of protective SARS-CoV-2 CD4^+^ and CD8^+^ T cell epitopes and characterization of the full repertoire of CD4^+^ and CD8^+^ T cells that are associated with the “natural protective immunity” seen in asymptomatic (ASYMP) COVID-19 patients (39). The recent availability of comprehensive genomic datasets of SARS-CoV-2 has shifted the paradigm of vaccine development from virological to sequence-based approaches (40). Our recent genome-wide screening of the SARS-CoV-2 sequence using cohorts of SYMP and ASYMP COVID-19 patients identified several previously unknown CD4^+^ and CD8^+^ T cell epitopes that span a wide range of structural, and non-structural SARS-CoV-2 proteins. Previously, we characterized the phenotype and function of CD4^+^ and CD8^+^ T cell epitopes in ASYMP COVID-19 patients and “humanized” HLA-DR*0101/HLA-A*0201 double transgenic mice. The present study extended our previous report and implemented an innovative prime/pull vaccine strategy that increases the size and function of SARS-CoV-2-specific CD4^+^ and CD8^+^ T cells in infected lungs. Our study demonstrated for the first time a protective efficacy against SARS-CoV-2 infection, COVID-19-like symptoms, and death in a “humanized” HLA-DR*0101/HLA-A*0201/hACE2 triple transgenic mouse model. Thus, our novel prime/pull vaccination strategy has confirmed an innovative delivery system that is prime for clinical trial testing.

One of the advantages of ASYMP epitope-based prime/pull vaccines, as opposed to protein-based prime/pull vaccines, is the avoidance of SYMP epitopes that might inadvertently drive unwanted immuno-pathological responses ultimately contributing to the exacerbation of COVID-19 inflammatory disease in the lungs. Thus, the identification of human “SYMP” epitopes from the SARS-CoV-2 genome and their elimination from future COVID-19 vaccines would be beneficial, since including such “SYMP” epitopes may be harmful by exacerbating COVID-19 inflammation (41).

The repertoire of SARS-CoV-2-specific CD4^+^ and CD8^+^ T cells typically targets epitopes from only a fraction of the viral proteins (42). Those viral epitopes usually fall into a dominance hierarchy consisting of one or a few dominant epitopes and several other sub-dominant epitopes. Through the use of a genome-wide comprehensive approach, we previously identified several new human CD4^+^ and CD8^+^ T cell epitopes from SARS-CoV-2 proteins not previously considered for epitope-based vaccine candidates. The present study confirmed the protective efficacy of the immunodominant CD4^+^ and CD8^+^ T cell epitopes.

The inadequacy of many animal models of SARS-CoV-2 infection and immunity has made it challenging to explore the immune mechanisms that lead to protection against COVID-19 (43). One critical question is which animal model would be the most appropriate to mimic the immuno-pathological aspects of COVID-19 symptoms as occurs in humans? ACE-2 transgenic mice have been the animal models of choice for most COVID-19 immunologists and the results from ACE-2 transgenic mouse model have yielded tremendous insights into the protective mechanisms during primary acute infection (44–46). Characterization of the phenotype and function of protective SARS-CoV-2-specific memory CD4^+^ and CD8^+^ T cells in ACE-2 transgenic mice have been largely limited to T cells specific to the immunodominant SARS-CoV-2 mouse epitopes studied during acute infection (47, 48). Considering the wealth of data addressing the protective mechanisms of CD4^+^ and CD8^+^ T cells specific to mouse SARS-CoV-2 epitopes, it is surprising how few reports exist exploring the protective mechanisms of CD4^+^ and CD8^+^ T cells specific to human SARS-CoV-2 epitopes. The present study generated a novel “humanized” susceptible HLA-DR*0101/HLA-A*0201/hACE2 triple transgenic mouse model and validated the COVID-19 disease in the “humanized” HLA-DR*0101/HLA-A*0201/hACE2 triple transgenic mouse model to pre-clinically test the safety immunogenicity and protective efficacy of our multi-epitope//CXCL11 prime/pull Coronavirus mucosal vaccine bearing human B, CD4^+^ CD8^+^ cell epitopes, Our findings demonstrate that both virus replication in the lungs and COVID-19 symptoms can be induced in SARS-CoV-2-infected HLA-DR*0101/HLA-A*0201/hACE2 triple transgenic mice. Moreover, our “humanized” HLA-DR*0101/HLA-A*0201/hACE2 triple transgenic mice express the human HLA-A*0201 and HLA-DR*0101-molecules, instead of mouse MHC molecules. Thus, they develop “human-like” CD4^+^ and CD8^+^ T-cell responses to HLA-restricted epitopes. In addition, high numbers of functional CD4^+^ and CD8^+^ T_RM_ and T_EM_ cells were detected in the lungs and were associated with protection against virus replication and COVID-like symptoms. In our opinion, the HLA-DR*0101/HLA-A*0201/hACE2 triple transgenic Tg mice are arguably the best available small animal model to study the role of HLA-restricted CD4^+^ and CD8^+^ T-cells specific to human SARS-CoV-2 epitopes in protection against virus replication and COVID-like symptoms.

The lung-tropic AAV9 vector used in our prime/pull vaccine has rapidly moved to the forefront of human therapies in the past few years (49). The AAV9 vector was not directly involved in reduced SARS-CoV-2 infection, COVID-19-like symptoms, and death, as seen in the AAV9-CXCL11 prime/pull vaccine since no protection, was observed in mice that received an empty AAV9 vector alone (not shown). Several naturally occurring, tissue-specific AAV serotypes have been isolated (50). We chose AAV9 because (*i*) it is a lung-tropic virus with the potential of persistent transgene expression in epithelial cells; (*ii*) it can superinfect SARS-CoV-2 infected cells; (*iii*) it can accommodate up to 4.7 kb of DNA; and (*iv*) it is nonpathogenic. Clinical trials using AAV vectors have shown only transient inflammation while demonstrating clinical benefits. No side effects were observed in SARS-CoV-2 infected mice during the 30 days of monitoring following treatment with the AAV9 vector. However, long-term monitoring of the lungs and brain for pathology associated with AAV9 will be necessary to ensure the vectors’ safety for eventual use in COVID-19 patients.

In conclusion, the present study: (*i*) validated previously unreported protective “ASYMP” epitopes that are potentially useful if included in a multi-epitope COVID-19 vaccine; (*ii*) characterizes the phenotype and the function of the protective CD4^+^ and CD8^+^ T cell sub-populations associated with immunologic control of SARS-CoV-2 infection, disease and deaths; and (*iii*) demonstrates that bolstering the number of functional SARS-COV-2-specific CD4^+^ and CD8^+^ T_EM_ and T_RM_ cells in the lungs through SARS-CoV-2 human epitopes/CXCL11-based prime/pull vaccine protected against SARS-CoV-2 infection, disease and death. Results from this pre-clinical research study should pave the way toward developing a novel clinical T-cell-based vaccine against COVID-19.

## MATERIALS AND METHODS

### Mice

Female K18-hACE2 transgenic mice (8-9 weeks old) were purchased from the Jackson Laboratory (Bar Harbor, ME). K18-hACE2 mice breeding was conducted in the UCI animal facility where female mice were used at 8-9 weeks. In addition, female HLA-DR*0101/HLA-A*0201/hACE2 triple transgenic mice (8-9 weeks old) were used. The HLA-DR*0101/HLA-A*0201/hACE2 triple transgenic mouse colony was established here at the UCI by cross-breeding K18-hACE2 mice (51) with double transgenic HLA-DR*0101/HLA-A*0201 mice (17). The animal studies were performed at the University of California Irvine and adhered to the Guide for the Care and Use of Laboratory Animals published by the US National Institute of Health. All animal experiments were performed under the approved IACUC protocol # AUP-22-086.

### Immunization and CXC chemokine treatment

Groups of (8-9 weeks old) female HLA-DR*0101/HLA-A*0201/hACE2 triple transgenic mice were immunized intranasally on day 0 with (2×10^10^ VP per mouse, *n* = 28) of a multi-epitope Coronavirus vaccine (CoV Vacc), consisting of highly conserved and immunogenic 16 CD8^+^ T cell epitopes, 6 CD4^+^ T cell epitopes, and 9 B cell epitopes. As a negative control, mice received sterile PBS (mock). Mice were treated intranasally with CXCL-9, CXCL-10, and CXCL-11 (2.4 μg in 20 μl of sterile PBS/mice). Recombinant murine MIG (CXCL-9), IP-10 (CXCL-10) and I-TAC (CXCL-11) were obtained (PEPROTECH, USA)

### SARS-CoV-2 infection

K18-hACE2 mice were infected intranasally with 1 x 10^4^ pfu of the SARS-CoV-2-USA-WA1/2020 variant in 20 μl sterile PBS. Following genital infection, mice were monitored daily for disease progression. Next, mice were treated intranasally with 3 μg of the CXCL-9, CXCL-10, and CXCL-11 diluted in sterile PBS. The mock group was treated with sterile PBS. The CXCL chemokine treatment was given on days 3, 5, 7, and 9 (p.i.). Mice were monitored daily for weight loss and survival until day 14 p.i. on which they were euthanized.

HLA-DR*0101/HLA-A*0201/hACE2 triple transgenic mice were Immunized with the Pan-CoV-Vacc (2×10^10^ VP per mouse, *n* = 28) or mock (*n* = 7) of untreated mice. The immunized mice were subsequently treated intranasally with 2.4 μg of CXCL-9 (*n* = 7), CXCL-10 (*n* = 7), and CXCL-11 (*n* = 7) on days 10, 12, 14, 22, 24, 26 post-immunization. On day 28 post-immunization, mice were intranasally infected with 1×10^4^ pfu of SARS-CoV-2 (USA-WA1/2020). Three mice groups were subsequently treated intranasally with CXCL-9 (*n* = 5), CXCL-10 (*n* = 5), and CXCL-11 (*n* = 5) on days 30, 32, and 34 post-immunization. On days, 4 and 8 oropharyngeal swabs were collected for virus RNA copy number measurement. The SARS-CoV-2 infected mice were monitored up to day 14 p.i., for disease, weight loss, and survival. On day 14 p.i. mice were euthanized, and lungs were collected for H & E staining.

### SARS-CoV-2 propagation and titration

VERO E6 cells isolated from the kidney of an African green monkey (ATCC, VA, USA) grown in Minimum Essential Medium Eagle with Earl’s salts and L-Glutamine (Corning, Manassas, VA) supplemented with 10% fetal bovine serum and 1% penicillin-streptomycin was used for virus propagation. SARS-CoV-2-USA-WA1/2020 variant was procured from Microbiologics (St. Cloud, MN). The virus SARS-CoV-2-USA-WA1/2020 provided by Microbiologics was originally isolated from an oropharyngeal swab sample of a patient with a respiratory illness who had recently returned from travel to the affected region of China and developed the clinical disease (COVID-19) in January 2020 in Washington, USA. As described previously, the SARS-CoV-2-USA-WA1/2020 variant was propagated in Calu3 cells (53). The virus was quantified by plaque assay in VERO E6.

### Flow cytometry

Single-cell suspensions from the mouse lungs after collagenase treatment (8mg/ml) for 1 hour were used for (fluorescence-activated cell sorter [FACS] buffer) staining. The following antibodies were used: anti-mouse CD4 (clone: RM4-4), CD8a (clone: 53-6.7), CD44 (clone: IM7), CD62L (clone: MEL-14), CD103 (clone: M290), and CD183 (CXCR3) (clone: CXCR3-173), (BD Biosciences). Surface staining was performed by adding mAbs against various cell markers to a total of 1 x 10^6^ cells in phosphate-buffered saline containing 1% FBS and 0.1% Sodium azide and left for 45 minutes at 4°C. Cells were washed three times with FACS buffer and fixed in PBS containing 2% paraformaldehyde (Sigma-Aldrich, St. Louis, MO).

### Immunohistochemistry

The HLA-DR*0101/HLA-A*0201/hACE2 triple transgenic mice lung sections were fixed in 4% PFA for 48 and then transferred to 70% ethanol. The tissue sections were then embedded in paraffin blocks and sectioned at 8 μm thickness. Slides were deparaffinized and rehydrated before hematoxylin and eosin (H&E) staining for routine immunopathology. Images were captured on the BZ-X710 All-in-One fluorescence microscope (Keyence).

### Virus titration in oropharyngeal swabs

Throat swabs were analyzed for SARS-CoV-2 specific RNA by qRT-PCR. As recommended by the CDC, we used *ORF1ab-specific* primers (Forward-5’-CCCTGTGGGTTTTACACTTAA-3’ and Reverse-5’-ACGATTGTGCATCAGCTGA-3’) and probe (6FAM-CCGTCTGCGGTATGTGGAAAGGTTATGG-BHQ) to detect the viral RNA level in lungs.

Briefly, 5 *m*l of the total nucleic acid eluate was added to a 20-*m*l total-volume reaction mixture (1x TaqPath 1-Step RT-qPCR Master Mix, CG [Thermo Fisher Scientific, Waltham, MA], with 0.9 *m*M each primer and 0.2 *m*M each probe). The RT-PCR was carried out using the ABI StepOnePlus thermocycler (Life Technologies, Grand Island, NY). When the Ct-value was relatively high (35 ≤ Ct < 40), the specimen was retested twice and considered positive if the Ct-value of any retest was less than 35.

### Statistical analysis

Data for each assay were compared by ANOVA and Student’s *t*-test using GraphPad Prism version 5 (La Jolla, CA). The physical estimation data were analyzed with the paired t-test using non-parametric Gaussian distribution based on the Wilcoxon matched-pairs signed rank test. As we previously described, differences between the groups were identified by ANOVA and multiple comparison procedures (12, 30). Data are expressed as the mean ± SD. Results were considered statistically significant at a *P* < 0.05.

## ACKNOWLEDGMENTS

This work is supported by the Fast-Grant PR12501 from Emergent Ventures, by a Gavin Herbert Eye Institute internal grant, by Public Health Service Research grants AI158060, AI150091, AI143348, AI147499, AI143326, AI138764, AI124911, and AI110902 from the National Institutes of Allergy and Infectious Diseases (NIAID) to LBM and by R43AI174383-01 to TechImmune, LLC.

