## Supplemental Figure 1 for "A Multi-Epitope/CXCL11 Prime/Pull Coronavirus Mucosal Vaccine Boosts the Frequency and the Function of Lung-Resident CD4^+^ and CD8^+^ Memory T Cells and Protects Against COVID-19-like Symptoms and Death Caused by SARS-CoV-2 infection"

*Supplemental Fig. S1*

**A**

**Anti-SARS-CoV-2 Spike-Specific IgG measured in sera from different groups of immunized and mock-immunized triple transgenic hACE2-HLA-A2/DR mice at day 27 post immunization.**

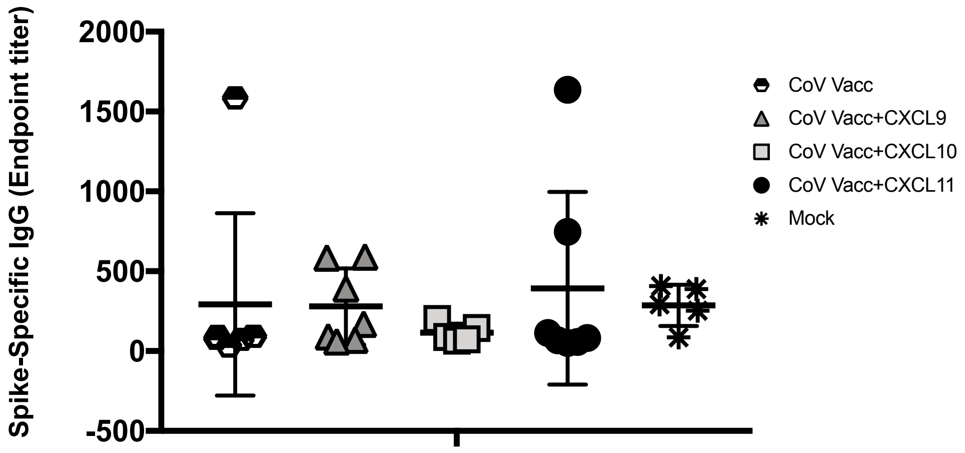


**B**

**SARS-CoV-2 Washington variant Neutralization by sera from different groups of immunized and mock-immunized triple transgenic hACE2-HLA-A2/DR mice at day 27 post immunization.**

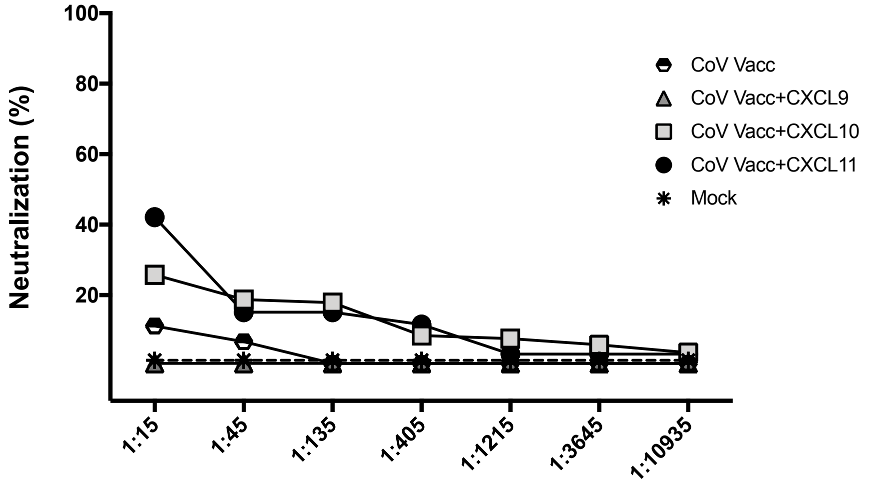


**C**

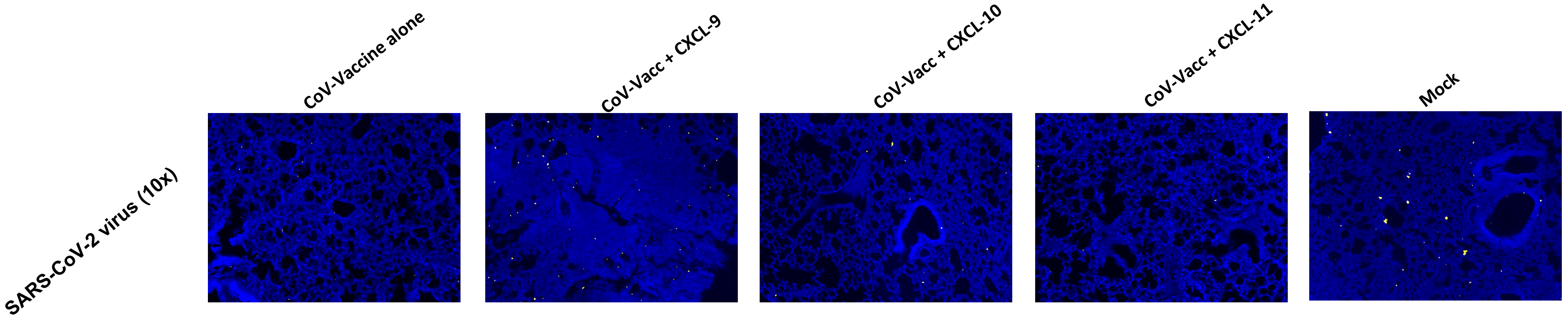


**Staining of triple transgenic hACE2-HLA-A2/DR mice lung tissue 14 days after exposure to SARS-CoV-2, for virus nucleoprotein (NP) in yellow (*n* = 3).**

*Supplemental Fig. S2*

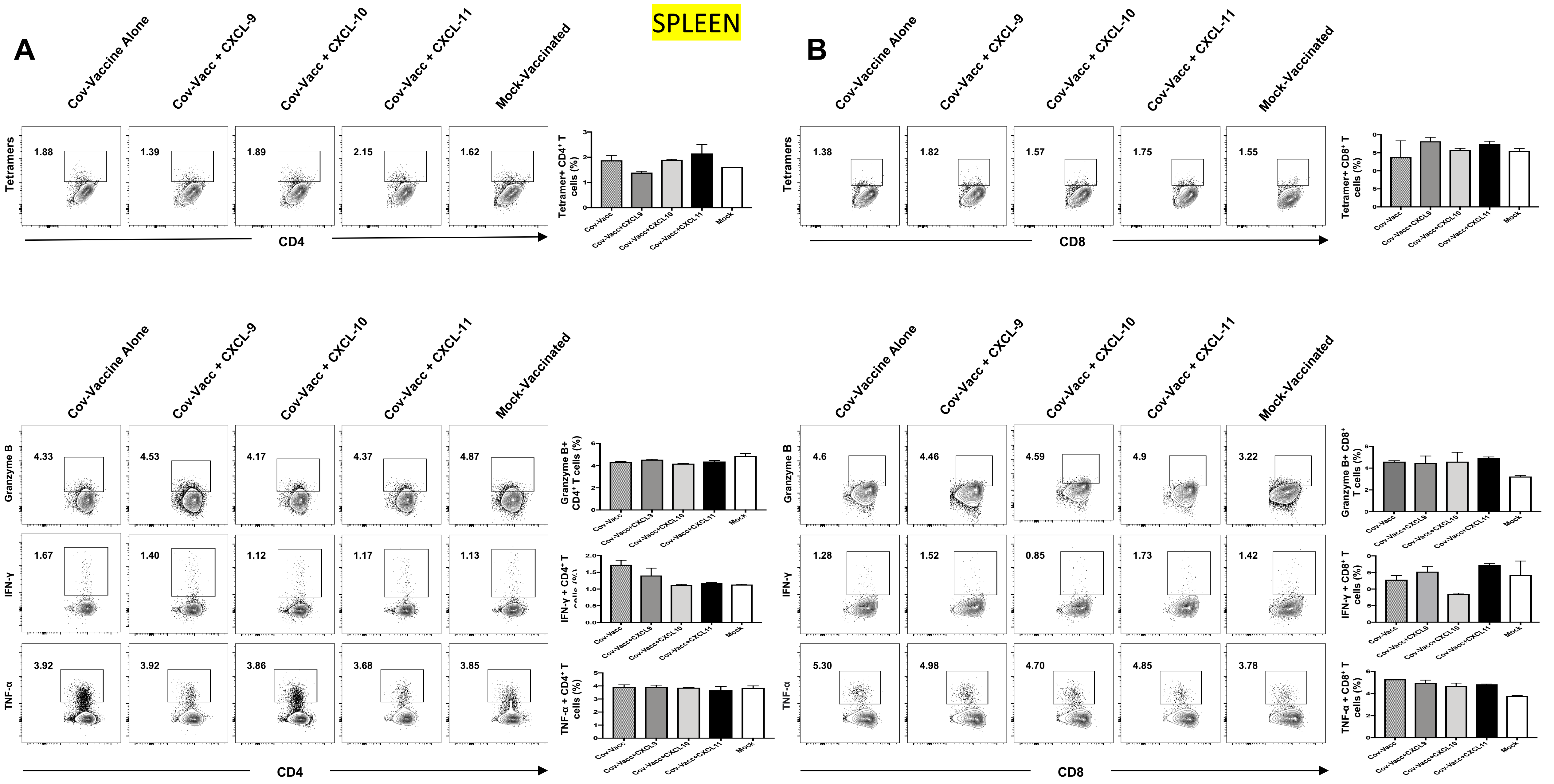
